# GGoutlieR: an R package to identify and visualize unusual geo-genetic patterns of biological samples

**DOI:** 10.1101/2023.04.06.535838

**Authors:** Che-Wei Chang, Karl Schmid

## Abstract

Landscape genomics is an emerging field of research that integrates genomic and environmental information to explore the drivers of evolution. Reliable data on the geographical origin of biological samples is a prerequisite for accurate landscape genomics studies. Traditionally, researchers discover potentially questionable samples using visualization-based tools. However, such approaches cannot handle large sample sizes due to overlapping data points on a graph and can hinder reproducible research. To address this shortcoming, we developed **G**eo-**G**enetic **outlier** (`GGoutlieR`), an R package of a heuristic framework for detecting and visualizing samples with unusual geo-genetic patterns. Outliers are identified by calculating empirical p-values for each sample, allowing users to identify them in data sets with thousands of samples. The package also provides a plotting function to display the geo-genetic patterns of outliers on a geographical map. GGoutlieR could significantly reduce the amount of data cleaning that researchers need to do before carrying out landscape genomics analyses.

## Statement of need

Landscape genomics is a thriving field in ecological conservation and evolutionary genetics (Aguirre-Liguori, Ramírez-Barahona, and Gaut 2021; Lasky, Josephs, and Morris 2023), providing insights into the links between genetic variation and environmental factors. This methodology requires reliable geographical and genomic information on biological samples. To determine whether data are reliable, researchers can examine associations between genetic similarities and the geographic origin of biological samples before proceeding with further studies. Under the assumption of isolation-by-distance, pairwise genetic similarities of samples are expected to decrease with increasing geographical distance between the sample origins. This assumption may be violated by long-distance migration or artificial factors such as human activity or data/sample management errors.

Visualization-based tools such as SPA (Yang et al. 2012), SpaceMix (Bradburd, Ralph, and Coop 2016), unPC (House and Hahn 2018) allow to identify samples with geo-genetic patterns that violate the isolation-by-distance assumption, but these tools do not provide statistics to robustly label outliers. Advances in genome sequencing technologies lead to much larger sample sizes, such as in geo-genetic analyses of genebank collections of rice (Gutaker et al. 2020; Wang et al. 2018), barley (Milner et al. 2019), wheat (Schulthess et al. 2022), soybean (Liu et al. 2020) and maize (Li et al. 2019). Visualization-based approaches may not be suitable to display unusual geo-genetic patterns in big datasets due to the large number of overlapping data points on a graph. To overcome this problem, we developed a heuristic statistical framework for detecting **G**eo-**G**enetic **outliers**, named GGoutlieR. Our GGoutlieR package computes empirical p-values for violation of the isolation-by-distance assumption for individual samples according to prior information on their geographic origin and genotyping data. This feature allows researchers to easily select outliers from thousands of samples for further investigation. In addition, `GGoutlieR` visualizes the geo-genetic patterns of outliers as a network on a geographical map, providing insights into the relationships between geography and genetic clusters.

## Algorithm of GGoutlieR

Under the isolation-by-distance assumption, the geographical origins of samples are predictable from their patterns of genetic variation (Battey, Ralph, and Kern 2020; Guillot et al. 2016), and vice versa. In this context, prediction models should result in large prediction errors for samples that violate the isolation-by-distance assumption. Based on this concept, we developed the GGoutlieR framework to model anomalous geo-genetic patterns.

Briefly, `GGoutlierR` uses *K* -nearest neighbour (KNN) regression to predict genetic components with the *K* nearest geographical neighbours, and also predicts in the opposite direction. Next, the prediction errors are transformed into distance-based (*D*) statistics and the optimal *K* is identified by minimising the sum of the *D* statistics. The *D* statistic is assumed to follow a gamma distribution with unknown parameters. An empirical gamma distribution is obtained as the null distribution by finding optimal parameters using maximum likelihood estimation. With the null gamma distribution, `GGoutlieR` tests the null hypothesis that the geogenetic pattern of a given sample is consistent with the isolation-by-distance assumption. Finally, p-values are calculated for each sample using the empirical null distribution and prediction error statistics. The details of the `GGoutlieR` framework are described step by step in the supplementary material (https://gitlab.com/kjschmid/ggoutlier/-/blob/master/paper/suppinfo.pdf).

## Example

### Outlier identification

For demonstration, we used the genotypic and passport data of the global barley landrace collection of 1,661 accessions from the IPK genebank (Milner et al. 2019; König et al. 2020). The full analysis of the barley dataset with `GGoutlieR` is available in the vignette (https://gitlab.com/kjschmid/ggoutlier/-/blob/master/vignettes/outlier_detection.pdf). Outliers were identified using the `ggoutlier` function. The function `summary_ggoutlier` was then used to obtain a summary table of outliers by taking the output of `ggoutlier`.

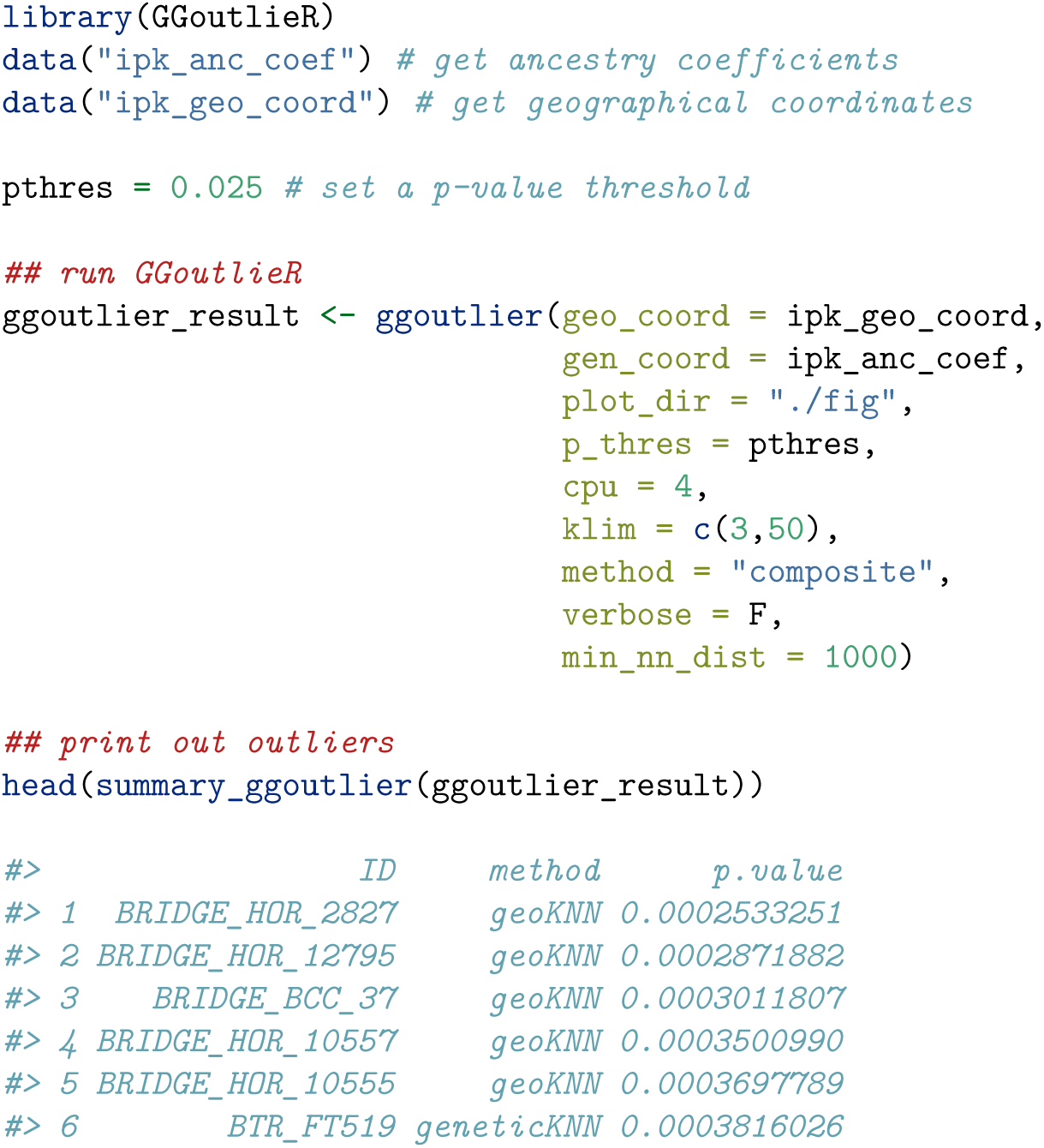

### Visualization of unusual geo-genetic patterns

The unusual geo-genetic patterns detected by GGoutlieR can be presented on a geographical map with the function `plot_ggoutlier` (Figure 1).

**Figure 1:**
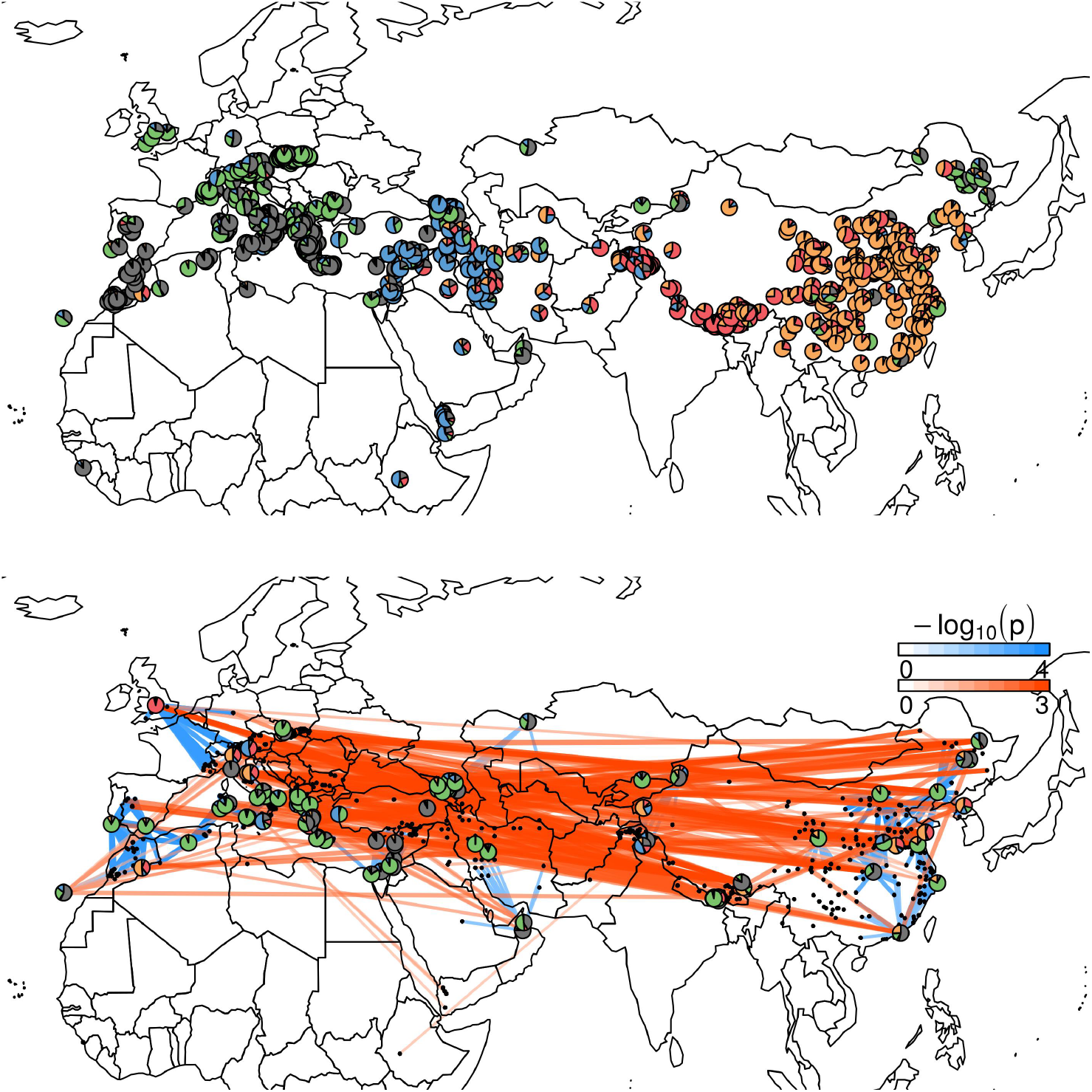
Visualization example of GGoutlieR with IPK barley landrace data. The red lines show the individual pairs with unusual genetic similarities across long geographical distances. The blue lines indicate the unusual genetic differences between geographical neighbors. Pie charts present the ancestry coefficients of outliers identified by GGoutlieR.

Moreover, the function `plot_ggoutlier` allows users to gain insight into outliers from a selected geographical region (Figure 2).

**Figure 2:**
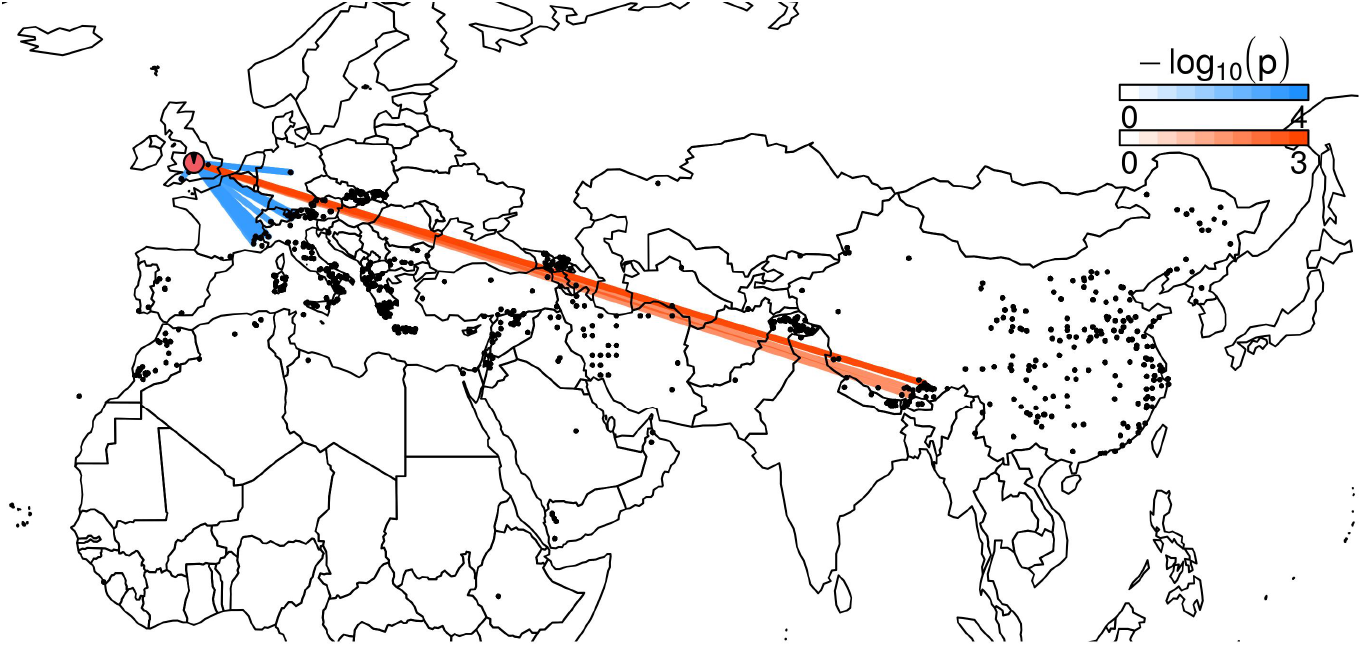
Visualization example of IPK barley landrace data with a highlight of samples from UK. The red lines show that the outliers in UK are genetically similar to accessions from Southern Tibet.

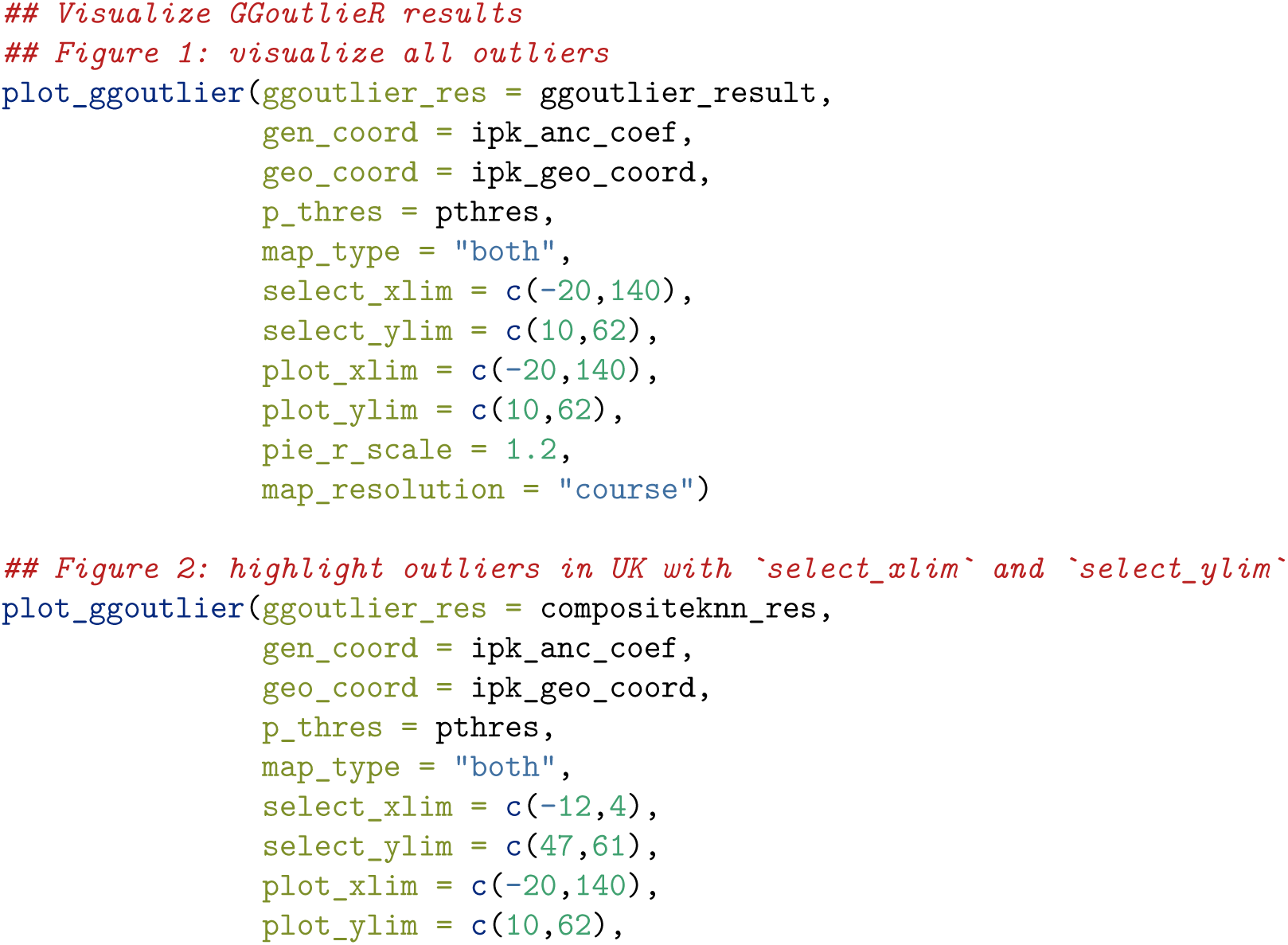
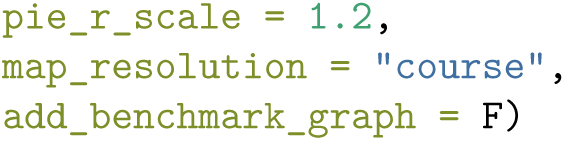

## Availability

The `GGoutlierR` package and vignette are available in our GitLab repository (https://gitlab.com/kjschmid/ggoutlier).

## Acknowledgement

We appreciate Dr. Martin Mascher and Max Haupt of Leibniz Institute of Plant Genetics and Crop Plant Research (IPK) for providing the raw VCF data of barley landraces that was used in the example. The project at Schmid lab was supported by the funds from the Federal Ministry of Food and Agriculture (BMEL) according to a decision of the parliament of the Federal Republic of Germany via the Federal Office for Agriculture and Food (BLE) under the Federal Programme for Ecological Farming and Other Forms of Sustainable Agriculture (Project number 2818202615). C.W.C was supported by the Study Abroad Fellowship from the Education Ministry of Taiwan (R.O.C.) (Project number 1100123625).

## References

Aguirre-Liguori, Jonas A, Santiago Ramírez-Barahona, and Brandon S Gaut. 2021. “The Evolutionary Genomics of Species’ Responses to Climate Change.” Nature Ecology & Evolution 5 (10): 1350–60. https://doi.org/https://doi.org/10.1038/s41559-021-01526-9.

Battey, Christopher J, Peter L Ralph, and Andrew D Kern. 2020. “Predicting Geographic Location from Genetic Variation with Deep Neural Networks.” eLife 9: e54507. https://doi.org/https://doi.org/10.7554/eLife.54507.

Bradburd, Gideon S, Peter L Ralph, and Graham M Coop. 2016. “A Spatial Framework for Understanding Population Structure and Admixture.” PLoS Genetics 12 (1): e1005703. https://doi.org/https://doi.org/10.1371/journal.pgen.1005703.

Guillot, Gilles, Hákon Jónsson, Antoine Hinge, Nabil Manchih, and Ludovic Orlando. 2016. “Accurate Continuous Geographic Assignment from Low-to High-Density SNP Data.” Bioinformatics 32 (7): 1106–8. https://doi.org/https://doi.org/10.1093/bioinformatics/btv703.

Gutaker, Rafal M, Simon C Groen, Emily S Bellis, Jae Y Choi, Inês S Pires, R Kyle Bocinsky, Emma R Slayton, et al. 2020. “Genomic History and Ecology of the Geographic Spread of Rice.” Nature Plants 6 (5): 492–502. https://doi.org/https://doi.org/10.1038/s41477-020-0659-6.

House, Geoffrey L, and Matthew W Hahn. 2018. “Evaluating Methods to Visualize Patterns of Genetic Differentiation on a Landscape.” Molecular Ecology Resources 18 (3): 448–60. https://doi.org/https://doi.org/10.1111/1755-0998.12747.

König, Patrick, Sebastian Beier, Martin Basterrechea, Danuta Schüler, Daniel Arend, Martin Mascher, Nils Stein, Uwe Scholz, and Matthias Lange. 2020. “BRIDGE–a Visual Analytics Web Tool for Barley Genebank Genomics.” Frontiers in Plant Science 11: 701. https://doi.org/https://doi.org/10.3389/fpls.2020.00701.

Lasky, Jesse R, Emily B Josephs, and Geoffrey P Morris. 2023. “Genotype– Environment Associations to Reveal the Molecular Basis of Environmental Adaptation.” The Plant Cell 35 (1): 125–38. https://doi.org/https://doi.org/10.1093/plcell/koac267.

Li, Jing, Guo-Bo Chen, Awais Rasheed, Delin Li, Kai Sonder, Cristian Zavala Espinosa, Jiankang Wang, et al. 2019. “Identifying Loci with Breeding Potential Across Temperate and Tropical Adaptation via EigenGWAS and EnvGWAS.” Molecular Ecology 28 (15): 3544–60. https://doi.org/https://doi.org/10.1111/mec.15169.

Liu, Yucheng, Huilong Du, Pengcheng Li, Yanting Shen, Hua Peng, Shulin Liu, Guo-An Zhou, et al. 2020. “Pan-Genome of Wild and Cultivated Soybeans.” Cell 182 (1): 162–76. https://doi.org/https://doi.org/10.1016/j.cell.2020.05.023.

Milner, Sara G, Matthias Jost, Shin Taketa, Elena Rey Mazón, Axel Himmelbach, Markus Oppermann, Stephan Weise, et al. 2019. “Genebank Genomics Highlights the Diversity of a Global Barley Collection.” Nature Genetics 51 (2): 319–26. https://doi.org/https://doi.org/10.1038/s41588-018-0266-x.

Schulthess, Albert W, Sandip M Kale, Fang Liu, Yusheng Zhao, Norman Philipp, Maximilian Rembe, Yong Jiang, et al. 2022. “Genomics-Informed Prebreeding Unlocks the Diversity in Genebanks for Wheat Improvement.” Nature Genetics 54 (10): 1544–52. https://doi.org/https://doi.org/10.1038/s41588-022-01189-7.

Wang, Wensheng, Ramil Mauleon, Zhiqiang Hu, Dmytro Chebotarov, Shuaishuai Tai, Zhichao Wu, Min Li, et al. 2018. “Genomic Variation in 3,010 Diverse Accessions of Asian Cultivated Rice.” Nature 557 (7703): 43–49. https://doi.org/https://doi.org/10.1038/s41586-018-0063-9.

Yang, Wen-Yun, John Novembre, Eleazar Eskin, and Eran Halperin. 2012. “A Model-Based Approach for Analysis of Spatial Structure in Genetic Data.” Nature Genetics 44 (6): 725–31. https://doi.org/https://doi.org/10.1038/ng.2285.

