## Supplementary material for "GGoutlieR: an R package to identify and visualize unusual geo-genetic patterns of biological samples"

### Methods

To facilitate the investigation of geo-genetic patterns, we introduced a heuristic framework, named **G**eo-**G**enetic **outlieR** (*GGoutlieR*). It quantifies the deviation from isolation-by-distance expectation on an individual basis, providing a data-driven baseline to identify outliers. The analytical framework is available in the R package *GGoutlieR*. Also, the package supports the visualization of unusual geo-genetic patterns. We described the *GGoutlieR* framework in detail below.

#### Overview

Under the isolation-by-distance (IBD) assumption, the association between geographical distances and genetic distances of individuals enables the prediction of geographical origins based on genetic variation and, reversely, the prediction of genetic components according to geographical origins. In both prediction scenarios, prediction errors of a given sample increase proportionally to the degree of deviation from the IBD expectation. Based on the concept mentioned above, we implemented the K-nearest neighbors (KNN) regression, a non-parametric and non-linear method, to characterize the geo-genetic relationship of each sample with its corresponding nearest neighbors. The prediction errors of KNN regression can be converted into distance-based statistics (hereafter named $D$ statistics), which are assumed to follow a Gamma distribution. Statistical tests are conducted based on the $D$ statistics to spot outlier samples deviating from the IBD expectation. The analytical framework is performed with the R function *ggoutlier* in the *GGoutlieR* package. Three approaches are available by setting the argument *method* in the function *ggoutlier*. We introduced them sequentially below.

#### Assumptions

To develop the *GGoutlieR* framework, we assume that IBD patterns are pervasive among samples, so the population genetic structure is generally accordant to geographical habitats. Let geographical distribution and genetic variation of samples be described with two coordinate systems, $S_{geo}$ and $S_{genetic}$. We assume that the coordinates of an individual in $S_{genetic}$ are predictable from its neighbors in $S_{geo}$ and vice versa if the IBD assumption holds.

We further assume that the prediction errors, defined as geographical distances between true and predicted coordinates in $S_{geo}$, follow a Gamma distribution with unknown parameters, written as $\Gamma_{geo}(\alpha, \beta)$. This assumption is made with the expectation of prediction errors approximating zero under IBD. Similarly, the mean of squared prediction errors of coordinates in $S_{genetic}$ is assumed to follow $\Gamma_{genetic}(\alpha, \beta)$.

#### Geo-genetic outlier detection with K nearest neighbors

##### Definition of coordinate system $S_{geo}$ and $S_{genetic}$

The sample coordinates in $S_{geo}$ are defined as geographical coordinates of collection sites with decimal degree format. For $S_{genetic}$, we used ancestry coefficients (Pritchard et al. 2000) to represent samples' coordinates for the empirical applications because ancestry coefficients are more interpretable and easier to visualize on a geographical map than principal component values. We regard a matrix of ancestry coefficients ($Q_{N \times F}$) estimated based on $F$ ancestral populations as the coordinates of $N$ samples distributing in a space $S_{genetic}$ with $F$ dimensions.

##### ***Approach 1***: outlier identification with geographical KNNs

The first approach in the *GGoutlieR* framework aims at identifying outliers that are genetically differentiated from their geographical KNNs.

**Step 1.** Compute pairwise geographical distance matrix.

**Step 2.** Find KNNs for each individual according to pairwise geographical distances with a given $K$. To avoid a divisor of zero in the equation 1 of **Step 3**, *GGoutlieR* will ignore neighbors within 100 meters by the default (controlled by the *min_nn_dist* argument). Otherwise, one unit of distance is added to the off-diagonal values of geographical distance matrix before searching KNNs if any pairwise distance is zero and *min_nn_dist* is set to zero.

**Step 3.** Predict $\hat{x}_{genetic,i,j}$ using a weighted KNN approach. $\hat{x}_{genetic,i,j}$ is the predicted coordinate of an individual $i$ in the dimension $j$ of $S_{genetic}$, where $i = \{1, 2, ..., N\}$ and $j = \{1, 2, ..., F\}$.

$N$ is the number of individuals and $F$ is the number of dimensions in $S_{genetic}$, i.e. the number of ancestral populations. The weight of the $k$ th nearest neighbor of an individual $i$ is computed as

$$w_{i,k} = \frac{\frac{1}{d_{i,k}}}{\sum_{k=1}^{K} \frac{1}{d_{i,k}}} \tag{1}$$

where $d_{i,k}$ is geographical distance between the individual $i$ and its $k$ th nearest neighbor. $\hat{x}_{genetic,i,j}$ is calculated as

$$\hat{x}_{genetic,i,j} = \frac{1}{K} \sum_{k=1}^{K} w_{i,k} \, x_{genetic,i,j,k} \tag{2}$$

where $K$ is a given number of nearest neighbors. The default of *GGoutlieR* searches the optimal $K$ with a range of values (see **Step 5.1**). $x_{genetic,i,j,k}$ is the coordinate of $k$ th neighbor of individual $i$ in the dimension $j$ of $S_{genetic}$.

**Step 4.** Compute mean of squared prediction errors as

$$D_{genetic,i} = \frac{1}{F} \sum_{j=1}^{f} \hat{\varepsilon}_{i,j}^{2} = \frac{1}{F} \sum_{j=1}^{f} (x_{genetic,i,j} - \hat{x}_{genetic,i,j})^2 \tag{3}$$

where $x_{genetic,i,j}$ and $\hat{x}_{genetic,i,j}$ are the true and predicted coordinates of the individual $i$ in the dimension $j$ of $S_{genetic}$, respectively.

**Step 5.1.** Search optimal number of nearest neighbors ($K$) by minimizing $\sum_{i=1}^{n} D_{genetic,i}$. The **Step 1-4** are repeated with a range of $K$ values (the default is from 3 to 50). As $D_{genetic}$ represents the size of prediction errors, we define optimal $K$ as the $K$ value resulting in the lowest $\sum_{i=1}^{n} D_{genetic,i}$.

**Step 5.2.** Repeat **Step 1-4** with the optimal $K$.

**Step 6.** Obtain an empirical null distribution $\Gamma_{genetic}(\alpha, \beta)$. $\alpha$ and $\beta$ are evaluated by maximum likelihood estimation.

**Step 7.** Test individuals with the empirical null distribution $\Gamma_{genetic}(\alpha, \beta)$ from **Step 6**. The null hypothesis is that a focal individual follows the IBD expectation, whereas the alternative hypothesis is that a focal individual is genetically differentiated from its $K$ geographically nearest neighbors.

Considering that a true outlier may induce the significance of its neighbors, we perform the test in a multi-stage manner. In each iteration, we drop the most significant individual and repeat the **Step 2-4** to exclude the influence from the most significant outlier. This procedure is repeated until no outlier is identified with a given significant level.

To use the genetic KNN approach, users have to set the argument *method = "geoKNN"* for the *ggoutlier* function.

##### ***Approach 2*: outlier identification with genetic KNNs**

The second approach of the *GGoutlieR* framework aims at identifying outliers that are geographically remote from genetically similar individuals, i.e. their corresponding KNNs in $S_{genetic}$. The rationale is similar to the first approach as described in the previous section.

**Step 1.** Compute pairwise Euclidean distances according to a given matrix of genetic components, i.e. ancestry coefficients. If any pairwise distance is zero, $10^{-6}$ is added to the off-diagonal values of the genetic distance matrix to avoid a divisor of zero in the equation 4 of **Step 3**. As an alternative option, *GGoutlieR* accepts a distance matrix given by users in this step if users prefer a customized calculation of individual-based genetic distances.

**Step 2.** Find KNNs for each individual according to pairwise genetic distances with a given $K$.

**Step 3.** Predict $\hat{x}_{geo,i,j}$ using a weighted KNN approach. $\hat{x}_{geo,i,j}$ is the predicted coordinate of an individual $i$ in the dimension $j$ of $S_{geo}$, where $i = \{1, 2, ..., N\}$ and $j = \{1, 2\}$. $N$ is the number of individuals and $j$ corresponds to longitude and latitude. The weight of the $k$ th nearest neighbor of an individual $i$ is computed as

$$w_{i,k} = \frac{\frac{1}{d_{i,k}^2}}{\sum_{k=1}^{K} \frac{1}{d_{i,k}^2}} \tag{4}$$

where $d_{i,k}$ is genetic distance between the individual $i$ and its $k$ th nearest neighbor computed in the **Step 1**. $\hat{x}_{geo,i,j}$ is calculated as

$$\hat{x}_{geo,i,j} = \frac{1}{K} \sum_{k=1}^{K} w_{i,k} \; x_{geo,i,j,k} \tag{5}$$

where $K$ is a given number of nearest neighbors. The default of *GGoutlieR* searches the optimal $K$ with a range of values (see **Step 5.1**). $x_{geo,i,j,k}$ is the coordinate of $k$ th neighbor of individual $i$ in the dimension $j$ of $S_{geo}$.

**Step 4.** Compute prediction errors as

$$D_{geo,i} = GeoDist(x_{geo,i}, \hat{x}_{geo,i}) \quad (6)$$

where $GeoDist(x_{geo,i}, \hat{x}_{geo,i})$ is the geographical distance between the true and predicted locations of the individual $i$, which is calculated with the *geosphere* package (Hijmans 2019).

**Step 5.1** Search optimal number of nearest neighbors ($K$) by minimizing $\sum_{i=1}^{n} D_{geo,i}$. The **Step 1 - 4** are repeated with a range of $K$ values. The $K$ value resulting in the lowest $\sum_{i=1}^{n} D_{geo,i}$ is considered as the optimal $K$ for the given data set. The default of *GGoutlieR* tests a range $K$ from 3 to 50.

**Step 5.2** Repeat **Step 1-4** with the optimal $K$.

**Step 6** Obtain an empirical null distribution $\Gamma_{geo}(\alpha, \beta)$. $\alpha$ and $\beta$ are identified by maximum likelihood estimation.

**Step 7** Test individuals with the empirical null distribution $\Gamma_{geo}(\alpha, \beta)$. The null hypothesis is that a focal individual follows the IBD expectation. The alternative hypothesis is that a focal individual is geographically remote from $K$ individuals that are genetically most similar to a focal individual. The test is carried out in a multi-stage manner as described in the **Step 7** of **Approach 1**.

To use the geographical KNN approach, users have to set the argument *method = "geneticKNN"* for the *ggoutlier* function.

##### ***Approach 3*: composite approach**

The geographical KNN and genetic KNN approach above attempt to identify geo-genetic outliers from different perspectives. To leverage both approaches, a composite method first carries out the **Step 1 - 6** of geographical KNN and genetic KNN approaches. Next, instead of doing multi-stage tests (**Step 7**) separately, the composite approach sequentially removes the most significant outlier among the results of two KNN approaches and then repeats the KNN searching and p-value computation to identify outliers with two KNN approaches. This iterative procedure continues until no new outlier raises with the given significant threshold.

To use the composite KNN approach, users have to set the argument *method = "composite"* for the *ggoutlier* function.

### References

Hijmans RJ (2019) geosphere: Spherical Trigonometry. URL https://CRAN.R-project.org/package=geosphere, r package version 1.5-10

Pritchard JK, Stephens M, Donnelly P (2000) Inference of population structure using multilocus genotype data. Genetics 155(2):945–959
